# GoPeaks: Histone Modification Peak Calling for CUT&Tag

**DOI:** 10.1101/2022.01.10.475735

**Authors:** William M Yashar, Garth Kong, Jake VanCampen, Brittany M Smith, Daniel J Coleman, Lucia Carbone, Galip Gürkan Yardimci, Julia E Maxson, Theodore P Braun

## Abstract

Genome-wide mapping of the histone modification landscape is critical to understanding transcriptional regulation. Cleavage Under Targets and Tagmentation (CUT&Tag) is a new method for profiling the localization of covalent histone modifications, offering improved sensitivity and decreased cost compared with Chromatin Immunoprecipitation Sequencing (ChIP-seq). Here, we present GoPeaks, a peak calling method specifically designed for histone modification CUT&Tag data. GoPeaks implements a Binomial distribution and stringent read count cut-off to nominate candidate genomic regions. We compared the performance of GoPeaks against commonly used peak calling algorithms to detect H3K4me3, H3K4me1, and H3K27Ac peaks from CUT&Tag data. These histone modifications display a range of peak profiles and are frequently used in epigenetic studies. We found GoPeaks robustly detects genome-wide histone modifications and, notably, identifies H3K27Ac with improved sensitivity compared to other standard peak calling algorithms.

## Introduction

Chemical modification of histone proteins is a key mechanism of transcriptional regulation. Histones package eukaryotic DNA into chromatin and control DNA conformation and organization. Post-translational modification of histone proteins alters chromatin structure and regulates the recruitment of nuclear proteins to DNA regulatory elements^1^. For example, trimethylation of histone 3 lysine 4 (H3K4me3) aids with the recruitment of positive transcriptional regulators to transcription start sites^2–4^. Similarly, acetylation of histone 3 lysine 27 (H3K27Ac) neutralizes the positive charge of the histone tail and loosens the interaction between histones and DNA, allowing access of transcription factors to DNA regulatory sequences^5^. Transcription factors are canonical regulators of transcription and binding of these factors is strongly correlated with histone modifications^6–9^. Large-scale studies have demonstrated that transcription factor-binding profiles can be used to predict histone modifications^10^. An understanding of genome-wide histone modifications is crucial to the understanding of transcriptional regulation.

Chromatin immunoprecipitation with sequencing (ChIP-seq)^2,11–13^, which couples antibodies that recognize histone modifications with next generation sequencing technology, has enabled genome-wide profiling of histone modifications. Although widely used for epigenetic profiling, ChIP-seq suffers from high background, artificial enrichment of highly expressed genes, and often requires prohibitively large number of cells per experiment^14,15^. Enzyme-tethering strategies including Cleavage Under Targets and Tagmentation (CUT&Tag) and Cleavage Under Targets & Release Using Nuclease (CUT&RUN) have been developed to overcome these issues and perform epigenetic profiling with a low number of cells and with minimal background^16^.

Epigenetic studies require mapping multiple histone modifications for a comprehensive understanding of transcriptional regulation. Detecting regions bound to lysine 4 residues on histone 3 that are mono-(H3K4me1) or trimethylated aids with the identification of promoters and enhancers, respectively, throughout the genome^17^. Co-localization of H3K4me1 and H3K4me3 with H3K27Ac is characteristic of activated DNA regulatory elements^18,19^. Genomic regions of modified histones in ChIP-seq and CUT&Tag are identified as stacks of aligned reads; such regions are called peaks. The peak profiles of common histone modifications are highly variable^2^, so algorithms that identify histone modification peaks need to robustly detect a range of peak profiles. While H3K4me3 peaks tend to be sharply localized, H3K4me1 peaks span a broader region^2^ (Figure 1). Moreover, H3K27Ac can mark large domains such as super-enhancers as well as discrete regions such as promoters, thus having both broad and narrow characteristics. In order to extract meaning for epigenetic studies reliant on histone modification CUT&Tag datasets, peak calling algorithms need to be flexible to identifying narrow and broad peak characteristics.

**Figure 1:**
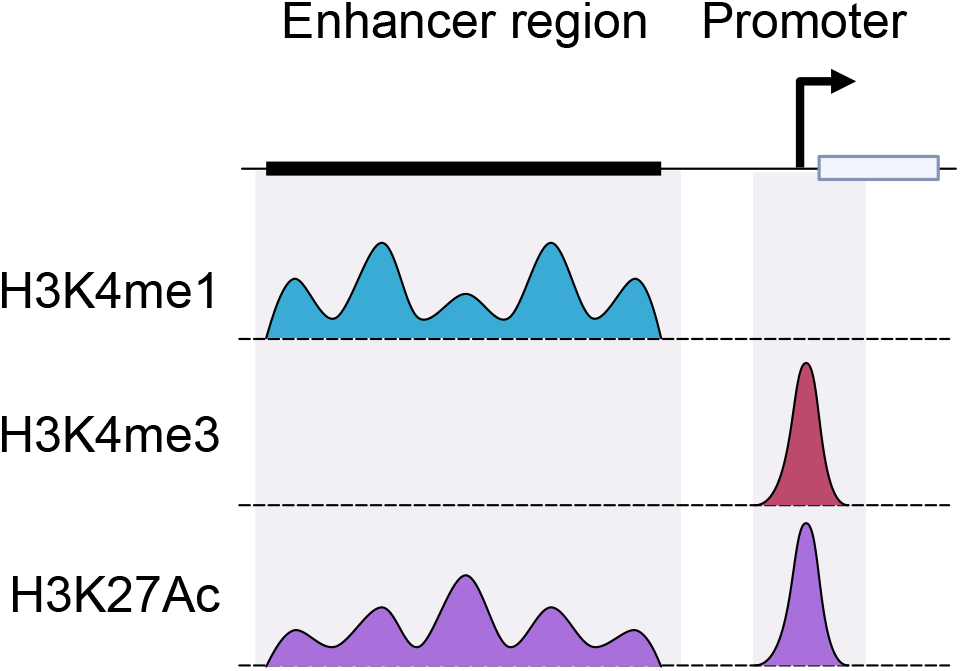
Histone modifications exhibit a range of peak profiles. Representative peak profiles for H3K4me1, H3K3me3, and H3K27Ac histone modifications.

Peak calling algorithms have been developed to not only identify genome-wide enrichment of aligned reads, but also to distinguish peaks of modified histones from noise and artifacts. Model-based Analysis of ChIP-Seq (MACS2)^20^, a widely-used peak calling algorithm for ChIP-seq, deploys a Poisson distribution to evaluate the likelihood that the proportion of aligned reads in a given region is statistically significant. However, MACS2 and other ChIP-seq peak calling methods are designed to address the high rate of background in ChIP-seq^21^ and are vulnerable to mistaking background signal as peaks particularly when the background is low^22^. While Sparse Enrichment Analysis (SEACR)^22^ was specifically designed for CUT&RUN, which is also characterized by low background, SEACR preferentially identifies broad, high-count peaks. No peak calling algorithms have been designed to address the low background and peak profile variability that is characteristic of histone modification CUT&Tag data.

Here, we present GoPeaks, a peak calling algorithm designed for histone modification CUT&Tag data. We compared the performance of GoPeaks against other widely used peak calling algorithms to detect H3K4me3, H3K4me1, and H3K27Ac peaks from CUT&Tag data. We demonstrate that GoPeaks robustly detects genome-wide histone modifications and notably, identifies H3K27Ac with improved sensitivity compared to other peak callers.

## Results

### Peak Calling with a Binomial Distribution and a Minimum Count Threshold

GoPeaks performs genome-wide peak identification of histone modification binding from CUT&Tag data in five general steps (Figure 2a). First, GoPeaks bins the genome into small intervals. Users can control the width of each bin with the “step” parameter and the width of bin overlap with the “slide” parameter. GoPeaks then quantifies the number of aligned reads contained within each bin and uses a Binomial distribution to determine whether the counts within each bin is significantly different from the genome-wide distribution of aligned reads. Bins with a significantly large number of counts are retained (p-value less than 0.05 before Benjamini-Hochberg correction by default). Moreover, bins must have a minimum number of counts to be retained (default of 15). Finally, significant bins that contain the minimum number of counts are merged into peaks if they lie within 150 bp of each other, which can be adjusted with the “mdist” parameter.

**Figure 2:**
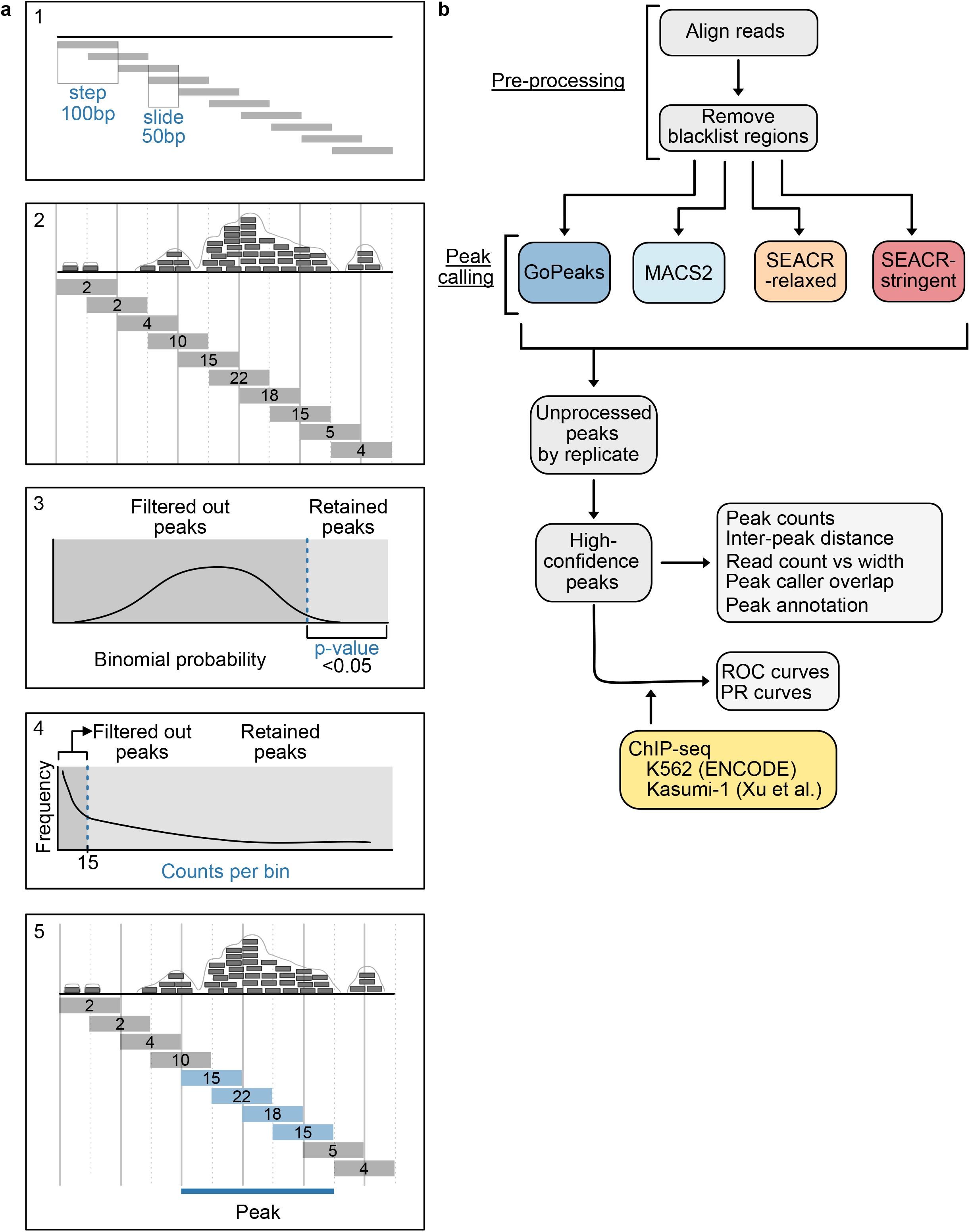
Overview of the GoPeaks methodology and benchmarking workflow. **a**. Five general steps of the GoPeaks peak calling methodology. Each subfigure (a1-a5) represents a separate step. (a1) Step indicates the bin width and slide, the width of the bin overlap. (a2) Counting the number of aligned reads per bin. (a3) Example of a Binomial probability test distribution and threshold to retain significantly different peaks. (a4) Filtering out bins with less than 15 counts. (a5) Retained bins within 150 bp are merged and identified as a peak. **b**. Schematic overview of the benchmarking workflow. All CUT&Tag datasets undergo the same pre-processing and are separately analyzed by the peak calling methods. The unprocessed peaks are extracted for sub-analyses. High-confidence peaks are those identified in both biological replicates. Generation of the receiver operator characteristic (ROC) and precision-recall (PR) curves require ChIP-seq standards.

We developed a computational workflow to compare the performance of GoPeaks against MACS2 and SEACR to identify histone modification peaks from CUT&Tag data (Figure 2b). Both SEACR threshold parameters that control peak selection, “SEACR-relaxed” and “SEACR-stringent”, were included in the workflow^22^. We evaluated each peak callers’ ability to identify peaks from CUT&Tag sequencing using publicly available H3K4me1 and H3K4me3 CUT&Tag data in K562 cells, a cell line model of blast-phase chronic myeloid leukemia (CML), and our H3K27Ac CUT&Tag data in Kasumi-1 cells, an acute myeloid leukemia (AML) cell line. Each CUT&Tag dataset was aligned to the GRCh38 genome and the ENCODE blacklist regions^23^ were removed. The unprocessed peaks and the high-confidence peaks, defined as statistically significant peaks present in at least two biological replicates, from each CUT&Tag dataset were used to quantify peak characteristics detected by each peak caller. We measured the sensitivity and specificity of each peak caller by their ability to recall peaks from publicly available ChIP-seq standards.

### Identification of Narrow H3K3me3 Peaks

To compare the performance of the peak calling algorithms on H3K4me3 marks, we assessed how many peaks each algorithm identified from the same CUT&Tag data on K562 cells. GoPeaks and MACS2 identified the greatest number of H3K4me3 peaks (Figure 3a). To assess the characteristics of the peaks called by each algorithm, we first calculated the average distance to the next nearest peak. This measurement indicates whether peak calling algorithms are splitting up peaks into smaller peaks and inflating the peak count total. We found that the peaks called by GoPeaks were similar distances apart as MACS2 and SEACR-relaxed (Figure 3b). MACS2, however, detected a small population of peaks less than 10^3^ bp apart. The peaks called by SEACR-stringent were noticeably farther apart than the other methods. To directly measure peak sizes identified by each peak calling method, we measured the number of counts in each peak and the peak width. We found that GoPeaks and MACS2 called peaks across a range of widths (Figure 3c). Both SEACR-relaxed and SEACR-stringent did not call any peaks with a width less than 1,000 bp, potentially missing or aggregating important regions. As an example, all peak callers recognized a peak overlapping the *CBX3* and *HNRNPA2B1* promoters approximately 8,500 bp wide (Figure 3d). Only GoPeaks identified a peak located at the promoter of *SNX10* (which has been implicated in pro-tumorigenic signaling^24^) nearly 1,450 bp wide. Together, these results demonstrate GoPeaks’ ability to identify H3K4me3 peaks across a range of sizes.

**Figure 3:**
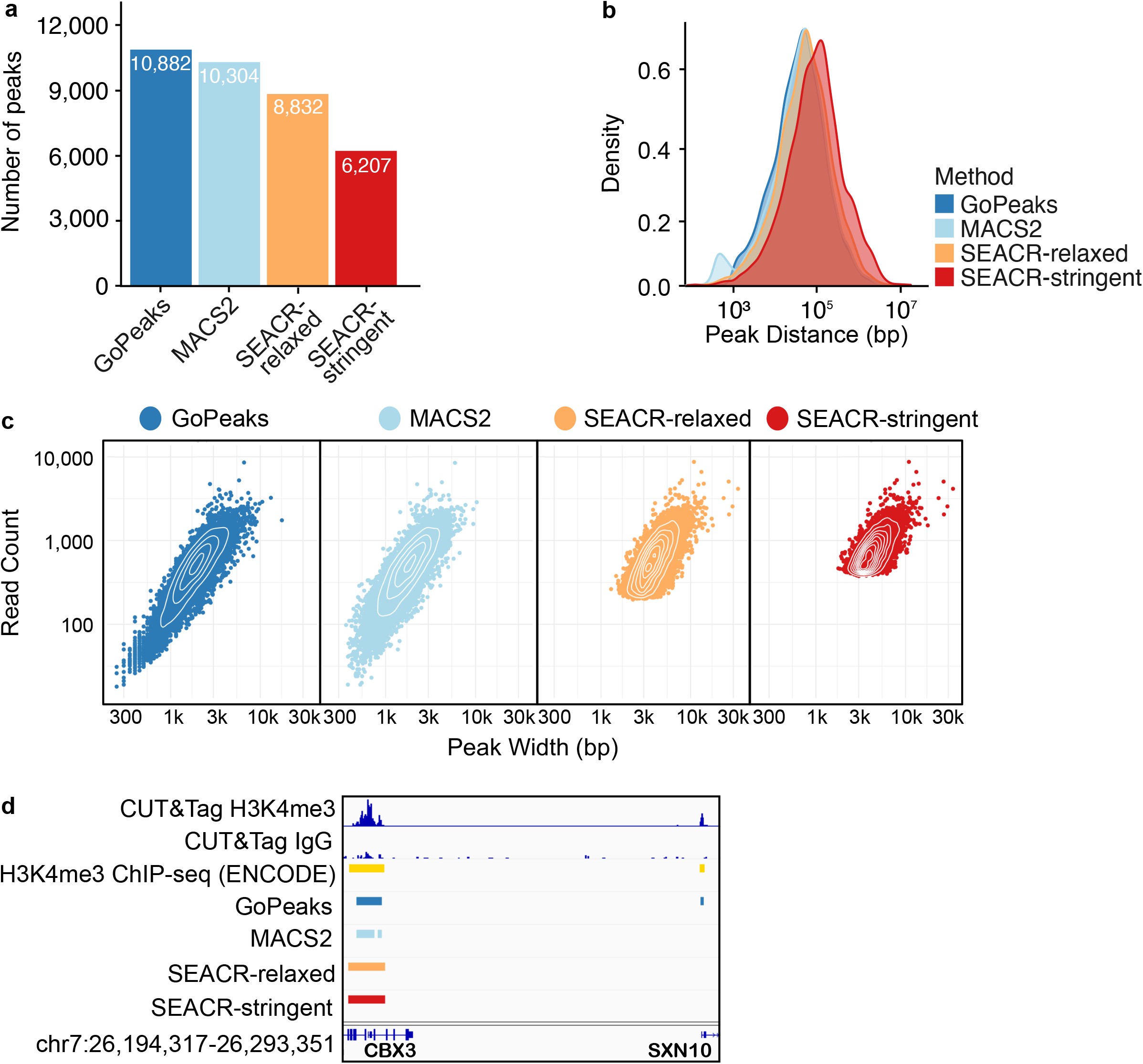
GoPeaks and MACS2 perform better than SEACR at identifying a range of H3K4me3 peak sizes. **a**. Number of high-confidence peaks identified from H3K4me3 CUT&Tag data in K562 cells per peak calling method. High-confidence peaks are those identified in two biological replicates. Colors indicate the peak calling method. **b**. Distribution of the distances to the next nearest peak. **c**. Distribution of read counts by peak width. Each dot represents the read count and peak width of a single detected peak. **d**. Example peaks at the *CBX3* and *SNX10* genes. IgG replicates are negative controls. Peak calls for each biological replicate are shown. Tracks are normalized by counts per million and are scaled to the range [0-5.10] by IGV. Tracks are depicted on the GRCh38 genome assembly.

### Sensitivity and Specificity of Detecting Narrow H3K4me3 Peaks

While both GoPeaks and MACS2 identify more H3K4me3 peaks than the SEACR peak calling methods, it is unclear whether some of these peaks may be false positives. To understand the sensitivity and specificity of each peak caller for H3K4me3 marks, we compared the peaks identified from publicly available K562 CUT&Tag data^16^ to those identified by ChIP-seq on the same cell line from the ENCODE Project^25^. We created receiver operating characteristic (ROC) curves, which maps the true positive rate, or recall, against the false positive rate. A true positive is defined as a peak identified in the K562 CUT&Tag data, which is also present in the ENCODE K562 ChIP-seq data. ROC curves along with precision-recall (PR) curves, which instead quantify the relationship between precision and recall, were used to characterize peak caller sensitivity and specificity. GoPeaks and MACS2 demonstrated a greater degree of peak recall than the SEACR methods for a given false positive rate (Figure 4a). GoPeaks and MACS2 had comparable area under the ROC curve (AUROC), which is a measurement of how well the peak callers detect CUT&Tag peaks that are also present in the ChIP-seq standard (Supplementary Figure 1). Every method demonstrated a similar ability to identify peaks with high precision across a range of recall values (Figure 4b). It is unlikely to observe perfect concordance due to the technical differences between the CUT&Tag and ChIP-seq assays. Indeed, given the sensitivity of CUT&Tag, it is probable that CUT&Tag will identify regions enriched with aligned reads that are not evident in ChIP-seq data.

**Figure 4:**
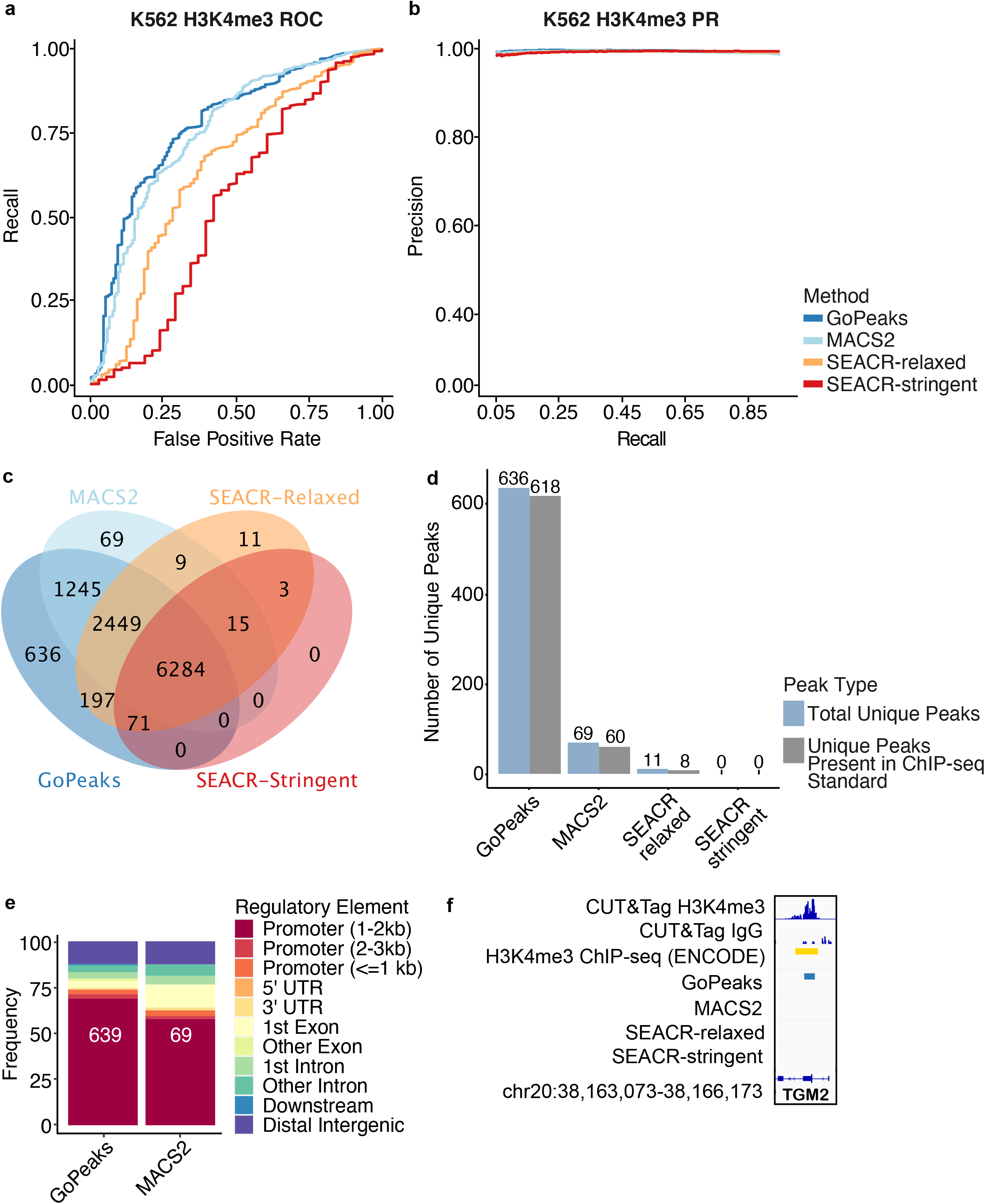
GoPeaks has a favorable specificity and sensitivity for narrow H3K4me3 CUT&Tag peaks. **a**. ROC curves quantifying the recall and false positive rates and **b**. PR curves quantifying the precision and recall rates of H3K4me3 CUT&Tag data from H3K4me3 ChIP-seq data. Both ChIP-seq and CUT&Tag datasets were generated in K562 cells. Colors indicate the peak calling method. **c**. Overlap of high-confidence peaks identified by each peak caller. High-confidence peaks are those identified in two biological replicates. **d**. Comparison of unique peaks that are identified by each peak calling algorithm and are also present in the ChIP-seq standard. Each bar is labeled by the number of peaks it represents. Colors indicate the peak type. **e**. Annotation of unique peaks identified by each peak caller. Colors indicate the putative region. Downstream is at least 300 bp towards 3’ end of DNA strand. **f**. Example peaks at the *TGM2* gene. IgG replicates are the negative controls. Peak calls for each biological replicate are shown. Tracks are normalized by counts per million and are scaled to the range [0-1.46] by IGV. Tracks are depicted on the GRCh38 genome assembly. UTR = untranslated region.

GoPeaks’ high sensitivity and specificity is likely due in part to its ability to identify peaks not captured by the other peak calling algorithms. To assess what may have distinguished each peak callers’ performance, we studied the overlap of the high-confidence peaks or peaks detected in both replicates. GoPeaks and MACS2 identified most peaks detected by both SEACR-stringent and SEACR-relaxed (Figure 4c). However, GoPeaks called 636 peaks not identified by any other peak caller, whereas MACS2 only identified 69 unique peaks. These unique peaks likely contributed to GoPeaks’ high sensitivity and specificity as 97.2% (618) of GoPeaks’ unique peaks were also present in the ChIP-seq standard (Figure 4d). Since H3K4me3 peaks are associated with promoters^17^, we annotated each peak set to the nearest gene feature. The unique peaks identified by GoPeaks and MACS2 were mostly associated with promoters (73.7% and 62.5%, respectfully), consistent with the established biology of H3K4me3 (Figure 4e). SEACR-relaxed and SEACR-stringent did not identify enough unique peaks to be included in the analysis. Notably, GoPeaks detected a peak in both CUT&Tag replicates located at the promoter of *TGM2*^26^ (Figure 4f). Studies have shown that *TGM2* is an important mediator of cell growth and differentiation^27^. Collectively, these results reveal that GoPeaks has favorable operating characteristics for H3K4Me3 data when compared with high-quality ChIP-seq, enabling the identification of an increased number of true positive peaks with a minimal false positive rate and at high precision.

### Sensitivity and Specificity of Detecting Broad H3K4me1 Peaks

While GoPeaks was highly sensitive and specific to narrow H3K4me3 peaks, we wanted to evaluate its performance to detecting broad H3K4me1 peaks. To compare the performance of each peak caller to detect H3K4me1 CUT&Tag peaks from K562 cells^16^, we measured their sensitivity and specificity against ENCODE H3K4me1 ChIP-seq on the same cell line^25^. Each peak caller identified peaks from publicly available H3K4me1 CUT&Tag sequencing in K562 cells. Overall, GoPeaks demonstrated comparable sensitivity and specificity across both H3K4me1 replicates (Figure 5a, b; Supplementary Figure 2a).

**Figure 5:**
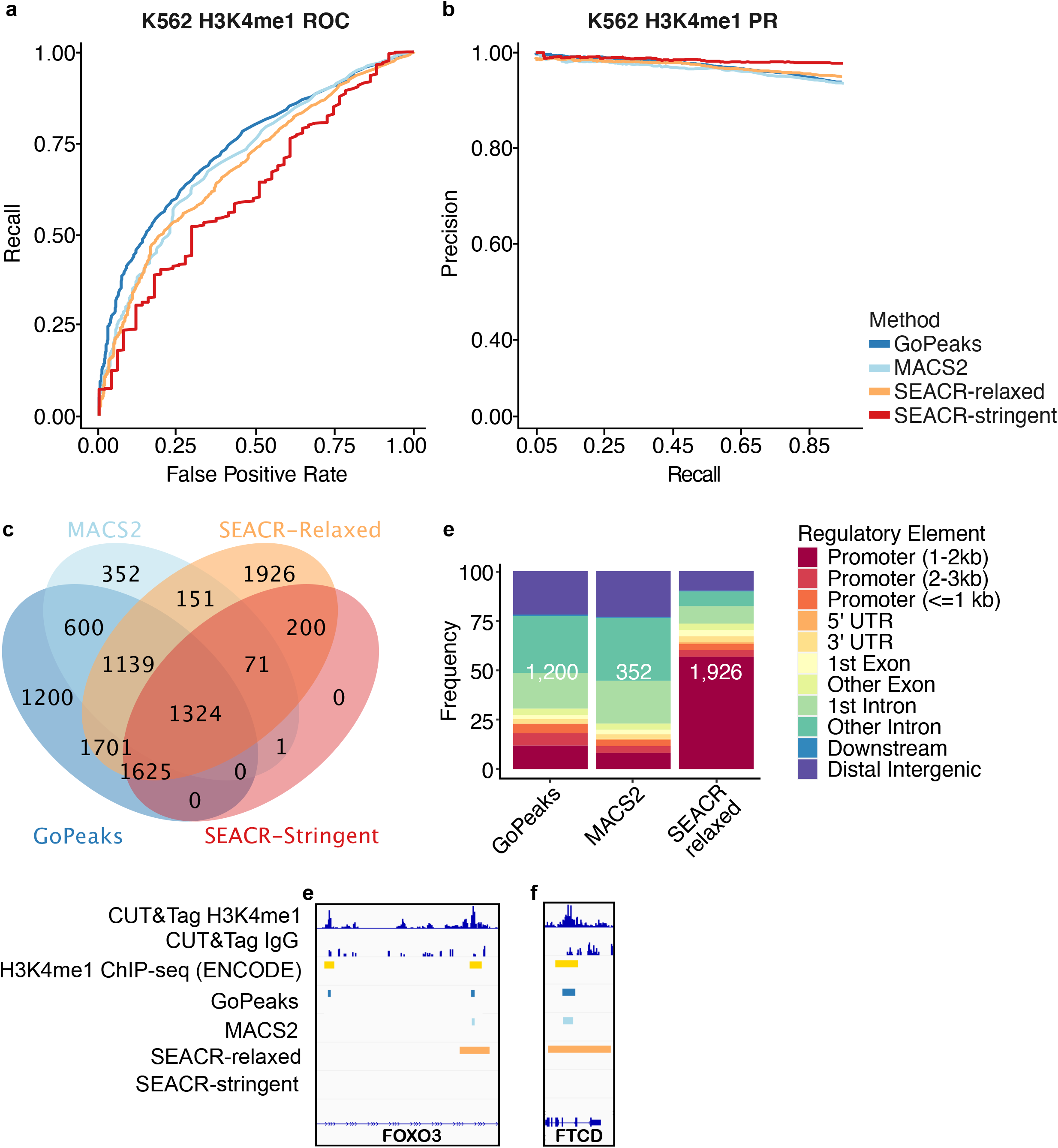
GoPeaks has a favorable specificity and sensitivity for broad H3K4Me1 CUT&Tag peaks. **a**. ROC curves quantifying the recall and false positive rates and **b**. PR curves quantifying the precision and recall rates of H3K4me1 CUT&Tag data from H3K4me1 ChIP-seq data. Both ChIP-seq and CUT&Tag datasets were generated in K562 cells. Colors indicate the peak calling method. **c**. Overlap of high-confidence peaks identified by each peak caller. High-confidence peaks are those identified in two biological replicates. **d**. Annotation of unique peaks identified by each peak caller. Colors indicate the putative region. Downstream is at least 300 bp towards 3’ end of DNA strand. Example peaks at the **e**. *FOXO3* gene and **f**. *FTCD* genes. IgG replicates are the negative controls. Peak calls for each biological replicate are shown. Tracks are normalized by counts per million and are scaled to the range [0-1.31] for D and [0-1.33] for E by IGV. Tracks are depicted on the GRCh38 genome assembly. UTR = untranslated region.

GoPeaks’ enhanced sensitivity and specificity may be due to its ability to detect H3K4me1 marks in intronic and intergenic regions. We therefore evaluated the overlap of the high-confidence peaks identified by each peak calling method. SEACR-relaxed identified the greatest number of unique H3K4me1 peaks among the four peak calling algorithms (Figure 5c). GoPeaks still identified 1,200 unique peaks, 85.7% (1,028) of which were also present in the ChIP-seq standard (Supplementary Figure 2b). To confirm the putative regions associated with the H3K4me1 peaks, we annotated each peak set to the nearest gene feature. The GoPeaks and MACS2 unique H3K4me1 peaks were primarily associated with intronic and intergenic regions (69.4% and 77.0%, respectively) whereas SEACR-relaxed were mostly associated with promoters (63.1%; Figure 5d). While H3K4me1 is found at active promoters, it displays the greatest enrichment at enhancers^2^. As an example, GoPeaks was able to identify a unique peak in the intronic region of *FOXO3*^*26*^, a family of proteins that play key roles in inducing mRNA expression of target genes involved in energy metabolism, apoptosis, cell cycle, DNA repair, cell death, and oxidative stress response^28–31^ (Figure 5e). Upstream of this unique peak, GoPeaks demonstrated its ability to detect the center of another H3K4me1 peaks. Both SEACR methods, on the other hand, called regions with widths greater than region annotated by the ChIP-seq standard. In regions where the intronic region is much smaller, like in *FTCD*, resolving the center of peaks is crucial (Figure 5f). Both SEACR methods identify genomic regions that include the promoter, exonic, and intronic regions of the *FTCD* gene, in contrast to what is detected by the ChIP-seq standard as well as GoPeaks. Together, these findings reveal that GoPeaks has favorable operating characteristics while simultaneously calling sufficiently narrow peaks to separate promoter and non-promoter regulatory regions.

### Sensitivity and Specificity of Detecting Broad & Narrow H3K27Ac Peaks

H3K27Ac marks are crucial for defining active regulatory elements^18,19^ and can have characteristics of broad and narrow peaks. To evaluate the performance of GoPeaks on H3K27Ac, we performed CUT&Tag sequencing for H3K27Ac on Kasumi-1 cells. We again measured the number and characteristic of peaks called in CUT&Tag data as compared to ChIP-seq. Since ENCODE does not have H3K27Ac ChIP-seq data for Kasumi-1 cells, we used the consensus of published H3K27Ac ChIP-seq data on the same cell line^32^. While GoPeaks showed an enhanced ability to recall peaks across a range of false positive rates (Figure 6a; Supplementary Figure 3a), this may have been at the expense of its PR characteristics (Figure 6b). Since precision is dependent on the number of true positives identified by each caller, we measured the number of peaks identified by each method. Indeed, GoPeaks detected 9,843 high-confidence peaks, which is nearly 3,000 more peaks than what was detected by the closest peak caller (Figure 6c). GoPeaks identified 2,907 peaks that were not detected by any other peak caller, 1,103 of which were also present in the standard (Figure 6d; Supplementary Figure 3b). In fact, GoPeaks identified 69.7% of all peaks present in the ChIP-seq standard (Figure 6e). In contrast, SEACR-stringent, which demonstrated improved PR characteristics over the other peak calling methods identified the least number of peaks (3,743). SEACR-stringent did not detect any unique peaks and only identified 33.0% of the peaks present in the standard. Overall, GoPeaks identified a substantial number of high-quality H3K27Ac peaks with some trade-off to its PR characteristics.

**Figure 6:**
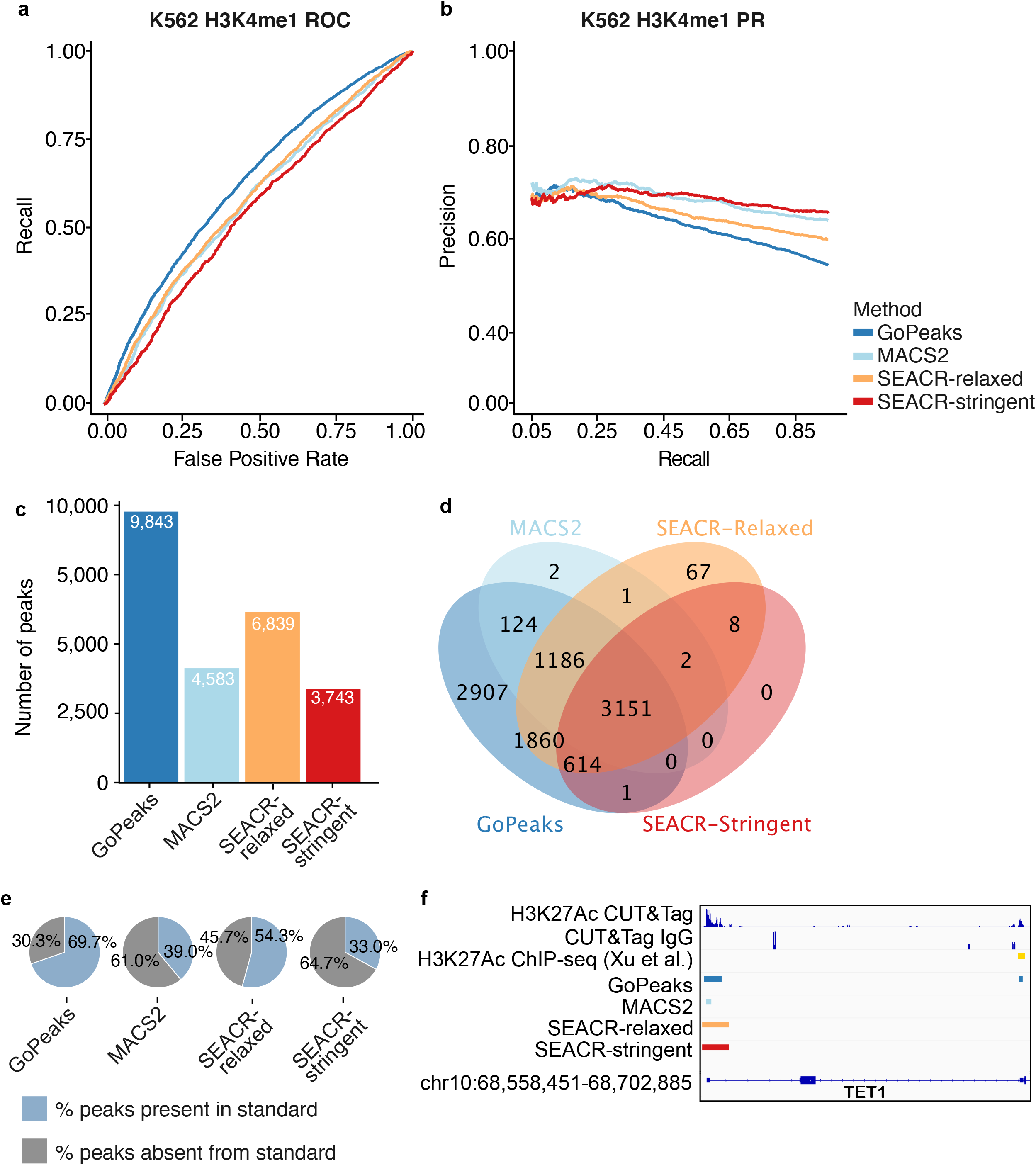
GoPeaks has higher specificity and sensitivity for H3K27Ac CUT&Tag peaks with broad and narrow peak shapes. **a**. ROC curves quantifying the recall and false positive rates and **b**. PR curves quantifying the precision and recall rates of H3K27Ac CUT&Tag data from H3K27Ac ChIP-seq data. Both ChIP-seq and CUT&Tag datasets were generated in Kasumi-1 cells. Colors indicate the peak calling method. **c**. Number of high-confidence peaks identified from H3K4me3 CUT&Tag data per peak calling method. High-confidence peaks are those identified in two biological replicates. **d**. Overlap of high-confidence peaks identified by each peak caller. **e**. Percent of total H3K27Ac ChIP-seq standard peaks that are identified by each peak caller. **f**. Example peaks near at the *TET1* gene. IgG replicates are the negative controls. Peak calls for each biological replicate are shown. Tracks are normalized by counts per million and are scaled to the range [0-4.26] by IGV. Tracks are depicted on the GRCh38 genome assembly.

GoPeaks was able to identify peaks across a range of widths, which is crucial for H3K27Ac peak detection (Supplementary Figure 3c). As an example, GoPeaks identified a peak with both narrow and broad characteristics at the promoter of *TET1*, an oncogene associated with leukemogenesis^30,31,32^ (Figure 6f). While GoPeaks and SEACR identified the whole peak, MACS2 only identified the narrow portion of the peak. Additionally, GoPeaks identified another narrow peak in an exonic region of *TET1*, which is also present in the ChIP-seq standard. Together, this data highlights GoPeaks’ dynamic range to identify both the narrow and broad peaks, which are characteristic of H3K27Ac marks.

## Discussion

GoPeaks was designed to address the low background and peak profile variability that defines histone modification CUT&Tag data. GoPeaks demonstrated a favorable ability to call peaks and were highly sensitive and specific to calling peaks across a range of histone modification CUT&Tag data. These results were particularly encouraging for H3K27Ac, which presents both broad and narrow peaks. Since H3K27Ac is a marker of active promoters and enhancers, it is crucial to pinpoint active regulatory non-coding elements.

MACS2 and SEACR both demonstrated biases towards the identification of narrow or broad peaks, respectively. MACS2 performed particularly well in analyzing H3K4me3 CUT&Tag data, in which peaks tend to be sharply localized. MACS2 identified a comparable amount of H3K4me3 CUT&Tag peaks as GoPeaks with similar operating characteristics. MACS2 was designed to identify narrow transcription factor peaks in ChIP-seq data^21^, so its bias for narrow peaks is unsurprising. SEACR, in contrast, demonstrated a favorable ability in detecting broad H3K4me1 peaks. SEACR-relaxed identified the most unique H3K4me1 peaks, but with comparable PR characteristics to GoPeaks. The reason for SEACR’s bias for broad marks may be due to its segmentation of the genome with contiguous, non-zero signal blocks^22^. SEACR empirically segments the genome into signal blocks with non-zero counts. Since the bin widths are not fixed, regions with low counts may be retained if they included in signal blocks that contain a true peak (Figure 5f). GoPeaks avoids this potential problem as each bin has a fixed width and is evaluated for significance before merging. GoPeaks’ simple but flexible framework is more amendable to the identification of both broad and narrow peaks, like those present in H3K27Ac data.

SEACR was more conservative in the identification of peaks across all marks. SEACR-stringent, in particular, consistently detected less peaks than the other methods. This strategy seems to be beneficial for SEACR’s PR characteristics, notably in the detection of H3K27Ac peaks, and may be more appropriate for researchers that have a low threshold identifying false positives. GoPeaks, on the other hand, may be best suited for researchers that are interested in discovering new peaks at the expense of some PR characteristics. However, GoPeaks largely performed at comparable PR characteristics and improved ROC characteristics over the other peak callers. Our analysis demonstrates GoPeaks detects a substantial number of high-quality histone modification peaks at high sensitivity and specificity.

There are important limitations to consider in this analysis. The performance of each peak calling method was only measured in three CUT&Tag histone modifications. Although the modifications studied are likely important for epigenetic studies, the peak profiles cover a broad range that will be likely encountered by other marks. We encourage users to test GoPeaks on other histone modification datasets. Additionally, we only tested these peak calling methods in two different cell lines. We cannot confidently rule out that GoPeaks may have a biological bias for K562 and Kasumi-1, although this is unlikely. The epigenetic profiles of K562 and Kasumi-1 cells are publicly available, which served as an important comparator for the ROC studies. Lastly, there were no CUT&Tag standards for the ROC studies. Although it would have been preferable to compare peaks between CUT&Tag datasets, the CUT&Tag technique is still new and few datasets exist in the public domain. However, our analysis indicates GoPeaks’ ability to extract biological meaning from CUT&Tag data. Overall, GoPeaks demonstrated to be a robust peak calling method across a range of histone modification CUT&Tag data.

## Methods

### GoPeaks Algorithm

GoPeaks detects peaks from aligned sequencing reads by first calculating the read coverage in sliding bins along the genome (*step* 100 bp and *slide* 50 bp by default). Read counts in a coverage bin are modeled by a Binomial distribution, described by two parameters “n” and “p”. The Binomial distribution *n* is equal to the total number of reads and *p* represents the probability of a bin containing *n* reads. *p* is estimated as the average read count in a non-zero coverage bin divided by the total number of reads. Coverage bins whose binomial probability of falling within this distribution below the given threshold are considered peaks (*p value* 0.05 before Benjamini-Hochberg correction by default). Modeling read counts using a Binomial distribution was originally inspired by algorithms from the Regulatory Genomics Toolbox^36^. Additionally, bins with fewer than “minreads” are filtered out (*minreads* 15 by default). Adjacent, overlapping, and bins within the distance defined by the parameter “mdist” are merged (*mdist* 150 bp by default). Finally, peaks are written to the output file if they are greater than the minimum width defined by the parameter “minwidth” (*minwidth* 150 bp by default).

#### Method Comparison Workflow

### Pre-Processing

K562 H3K4me3 (GEO accession GSM3536516), H3K4me1 (GEO accession GSM3536518), and IgG (GEO accession GSM3560264) CUT&Tag data from Kaya-Okur et al. 2019^16^ were downloaded through National Center for Biotechnology Information Gene Expression Omnibus (NCBI GEO)^37^. All CUT&Tag data was aligned to the GRCh38 genome with Bowtie2^13^ with the following options “--local --very-sensitive-local --no-unal --no-mixed --no-discordant --phred33 -I 10 -X 700”. The reads in the ENCODE GRCh38 blacklisted regions^23^ were removed prior to peak calling.

### Peak Calling

GoPeaks (v0.1.7) used the optional flag “-mdist 1000” to merge peaks within 1 kbp. MACS2^20^ (v 2.2.7.1) used the “--format BAMPE” flag with a genome size of 2.7e9 and the standard FDR threshold of 0.05. SEACR^22^ (v 1.3) used the “norm” flag when treatment and IgG samples were used in addition to using the relaxed and stringent mode. SEACR uses an empirical false discovery rate (FDR) calculated by quantifying the percentage of control signal blocks remaining out of the total above the threshold^22^.

### Post-Processing

After peaks were called for each method, high-confidence peaks were selected by taking the union of peaks that appear in at least two biological replicates within a study’s data set via a custom script. The purpose of finding high-confidence peaks is to reduce spurious peaks called in only one replicate and focus on peaks that consistently appear in multiple replicates. Intervene^39^ was used on the high-confidence peak sets to find common and exclusive peaks across peak callers.

### Peak Characterization

Peak counting was done in base R. ChIPseeker^40^ was used to annotate peaks to the nearest transcription start site. The read count at high-confidence peak intervals was tallied with BEDtools^41^ intersect –C to yield read depth density distributions, and peak-peak distances were calculated with GRanges^42^. Data cleaning and visualization were mainly facilitated using data.table and ggplot2^43^. Tracks were normalized by counts per million (CPM) and visualized using Integrative Genomics Viewer (IGV)^26^.

### Receiver Operating Characteristic and Precision-Recall Curves

In the receiver operating characteristic (ROC) and precision-recall (PR) analyses, the ranking metrics for each peak calling algorithm was counts at high-confidence peaks (obtained through BEDtools intersect -C). The outputs of MACS2, SEACR, and GoPeaks high-confidence peak counts were the input for ROC and PR analyses. These high-confidence counts were compared to publicly available ChIP-seq standards downloaded from the ENCODE portal^25,44^ and ChIP-Atlas^45^. K562 H3K4me3 (ENCODE ID ENCFF885FQN) and H3K4me1 (ENCODE ID ENCFF759NWD) ChIP-seq data was accessed from the ENCODE portal^25,44^. Kasumi-1 H3K27Ac (ChIP-Atlas SRX ID SRX4143063 and SRX4143067) ChIP-seq data^32^ was accessed from ChIP-Atlas^45^. The standards were filtered for peaks with log_10_(p-value) > 10 and adjacent peaks were merged if they were within 1 kb.

Custom scripts were used to threshold over unique values of each ranking metric to define predicted truth and false, which were intersected with the ChIP-seq standards to fill out the confusion matrix. True negatives are defined as peaks that did not meet the threshold for significance and were not annotated in the ChIP-seq standard. Secondary properties such as precision, recall, and FPR, were calculated, and ROC curves were made by plotting precision versus false positive rate. The area under the curve was approximated with Riemann Sums using trapezoids.

#### Cell Culture and CUT&Tag Methods

### Cell Lines

Kasumi-1 cells (ATCC) were cultured in RPMI (Gibco) supplemented with 20% fetal calf serum (FCS, HyClone), 2 mM GlutaMAX (Gibco), 100 units/mL Penicillin, and 100 ug/mL Streptomycin (Gibco). Cells were cultured at 5% CO_2_ and 37°C. Cell lines were tested monthly for mycoplasma contamination.

### CUT&Tag

Benchtop CUT&Tag was performed as previously described^8^. In brief, Kasumi-1 cells were counted, harvested, and centrifuged for 5 min at 300xg at room temperature. Cells were washed two times in 1.5 mL wash buffer (20□mM HEPES pH 7.5, 150□mM NaCl, 0.5□mM Spermidine, 1× Protease inhibitor cocktail). Concanavalin A magnetic coated beads (Bangs Laboratories) were activated in binding buffer by washing two times (20 mM HEPES pH 7.5, 10 mM KCl, 1 mM CaCl_2_, 1 mM MnCl_2_). Washed cells were separated into 100,000 cell aliquots and 10 ul of activated beads were added to each sample. Samples rotated at room temperature end over end for 7 minutes. Beads were separated with a magnetic and supernatant was removed. Primary antibody was diluted 1:50 in antibody buffer (20 mM HEPES pH 7.5, 150mM NaCl, 0.5 mM Spermidine, 1× Protease inhibitor cocktail, 0.05% digitonin, 2 mM EDTA, 0.1% BSA). The primary antibodies used were: H3K27Ac (ab4729, Abcam) and Normal Rabbit IgG (#2729, CST). Cells were incubated overnight at 4°C on a nutator. Primary antibody was replaced with a guinea-pig anti rabbit secondary antibody diluted to 1:100 in wash buffer (Antibodies Online). Samples were incubated for 45 minutes at room temperature on nutator. Secondary antibody was removed, and samples were washed 2X in dig-wash buffer (20□mM HEPES pH 7.5, 150□mM NaCl, 0.5□mM Spermidine, 1× Protease inhibitor cocktail, 0.05% Digitonin). pA-Tn5 transposase, prepared and loaded with adaptors as previously described^16^, was diluted 1:250 in dig-300 buffer (20 mM HEPES pH 7.5, 300 mM NaCl, 0.5 mM Spermidine, 1× Protease inhibitor cocktail, 0.01% digitonin) and added to samples. Samples incubated for 1 hour at room temperature on nutator. Samples were washed 2X with dig-300 buffer then resuspended in tagmentation buffer (dig-300 buffer with 10 mM MgCl_2_). Samples were incubated at 37°C for 1 hour. DNA was extracted with a DNA Clean & Concentrator-5 kit (ZYMO). Samples were amplified by PCR using custom Nextera primers at 400 nM and NEBNext HiFi 2x PCR Master Mix (New England Biolabs)^46^. PCR conditions were set to: 72°C for 5 minutes, 98°C for 30 seconds, 14-27 cycles of 98°C for 10 sec, 63°C for 10 sec, and 72°C for 1 minute. Libraries were purified with AMPure Beads (Beckman) and sequenced on a NextSeq 500 sequencer (Illumina) using 37 BP PE sequencing by Massive Parallel Sequencing Shared Resource at Oregon Health and Science University.

## Availability of data and materials

GoPeaks is free to use and is publicly accessible on GitHub: https://github.com/maxsonBraunLab/gopeaks. Custom scripts used to compare the peak calling algorithms are available in the gopeaks-compare repository: https://github.com/maxsonBraunLab/gopeaks-compare. The datasets supporting the conclusions of this article are available upon request.

## Acknowledgements

We thank the following OHSU core facilities for their assistance: Massive Parallel Sequencing Shared Resource, ExaCloud Cluster Computational Resource, and the Advanced Computing Center. Funding was provided by an American Society of Hematology Research Restart Award, an American Society of Hematology Scholar Award and 1 K08 CA245224 from NCI awarded to TPB. WMY was supported by NIH T32 GM109835: Medical Scientist Training Program of Oregon Health and Science University.

## Author contributions

BMS and DJC identified the need for a new peak caller. JV quantitatively described the problem and wrote GoPeaks. WMY, GLK, JV, BMS, and TPB designed the research plan. BMS performed sample and library preparation. GLK designed and executed the benchmarking workflow. WMY, GLK, JV, BMS, GGY, and TPB analyzed data. WMY wrote the manuscript. WMY, BMS, TPB, LC, GGY, and JEM reviewed and edited the manuscript. TPB and JEM acquired funding and supervised the project. All co-first authors may identify themselves as lead authors in their respective CVs.

---

**Supplementary Figure 1:**
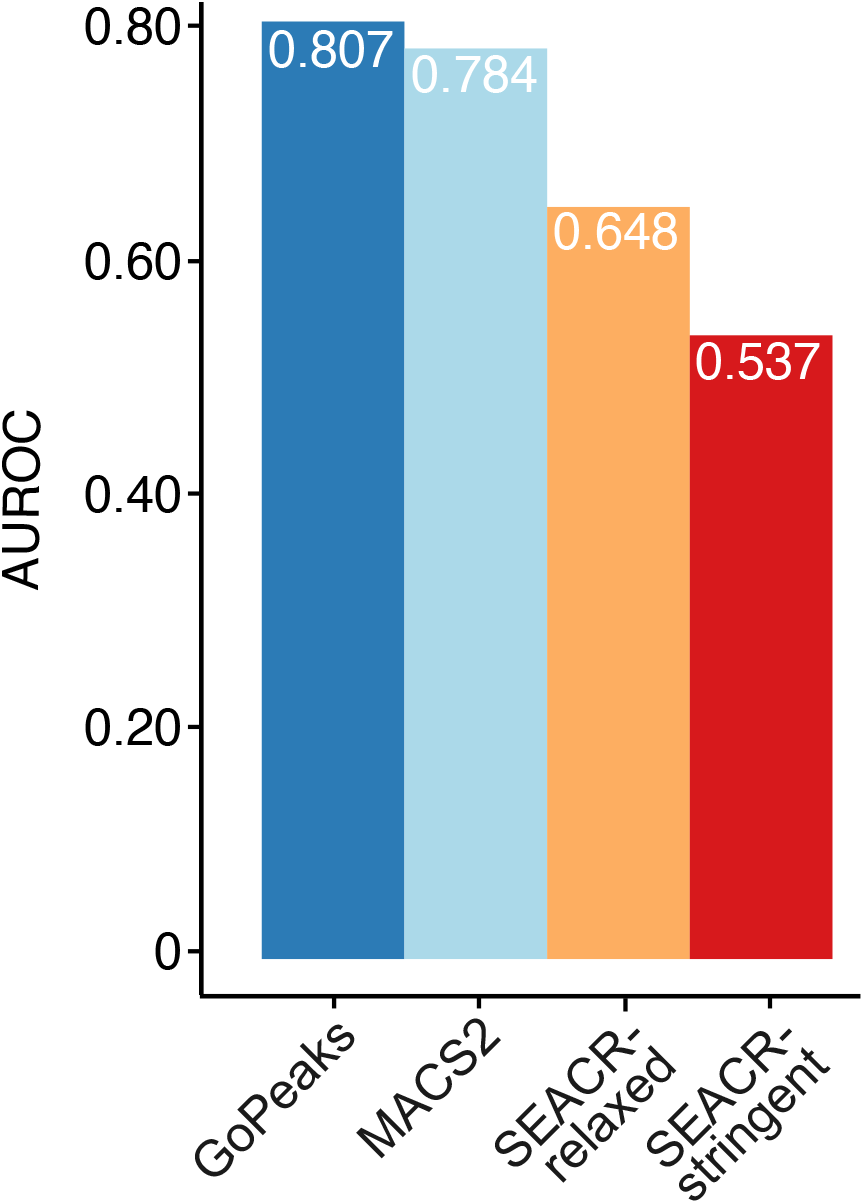
GoPeaks demonstrates comparable sensitivity and specificity in identifying H3K4me3 ChIP-seq standard peaks from CUT&Tag data. Area under the ROC curve (AUROC) for each peak calling method. Each bar is labeled by the AUROC value it represents. Colors indicate the peak calling method.

**Supplementary Figure 2:**
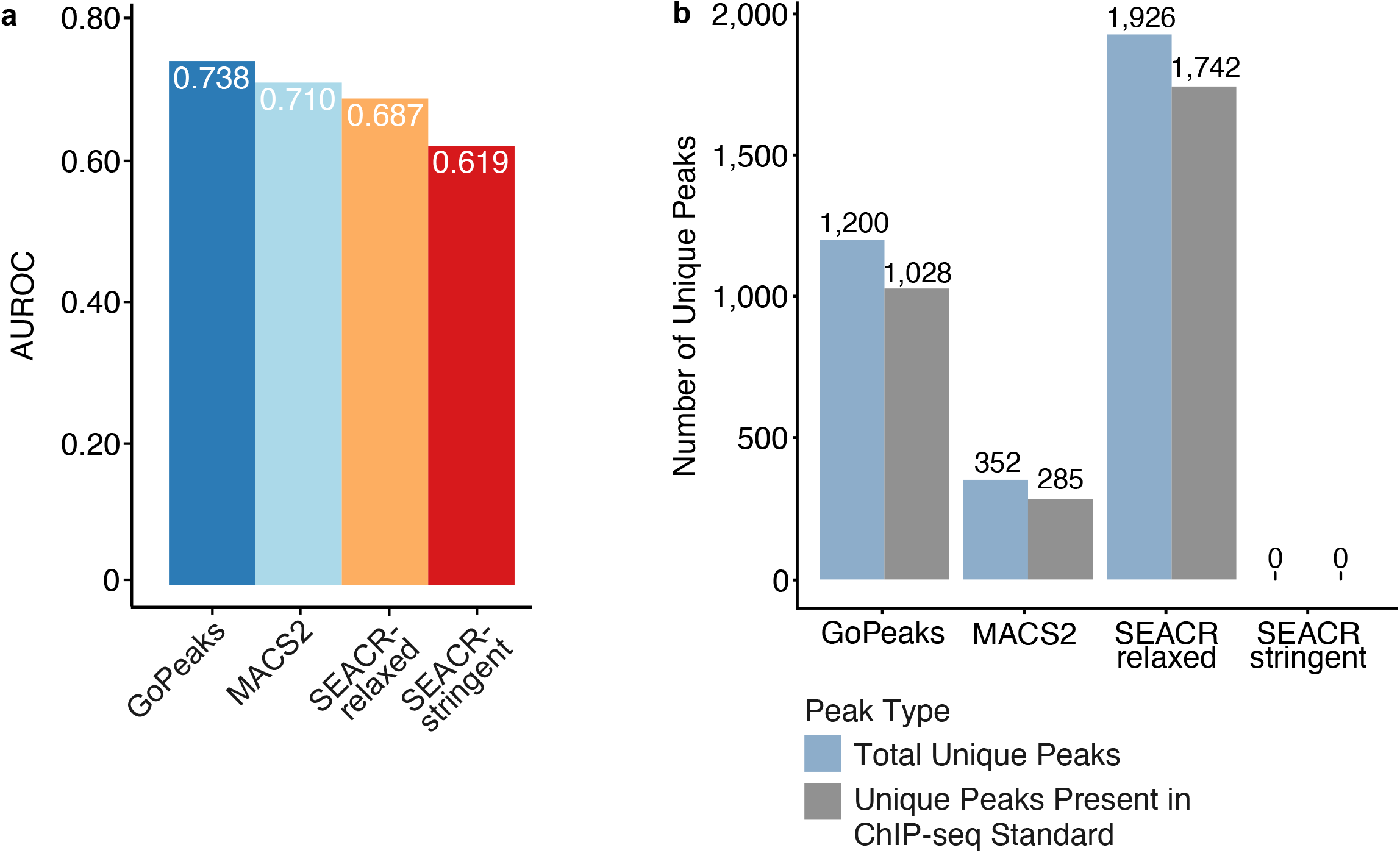
GoPeaks demonstrates comparable sensitivity and specificity in identifying H3K4me1 ChIP-seq standard peaks from CUT&Tag data. **a**. Area under the ROC curve (AUROC) for each peak calling method. Each bar is labeled by the AUROC value it represents. Colors indicate the peak calling method. **b**. Comparison of unique peaks that are identified by each peak calling algorithm and are also present in the ChIP-seq standard. Each bar is labeled by the number of peaks it represents. Colors indicate the peak type.

**Supplementary Figure 3:**
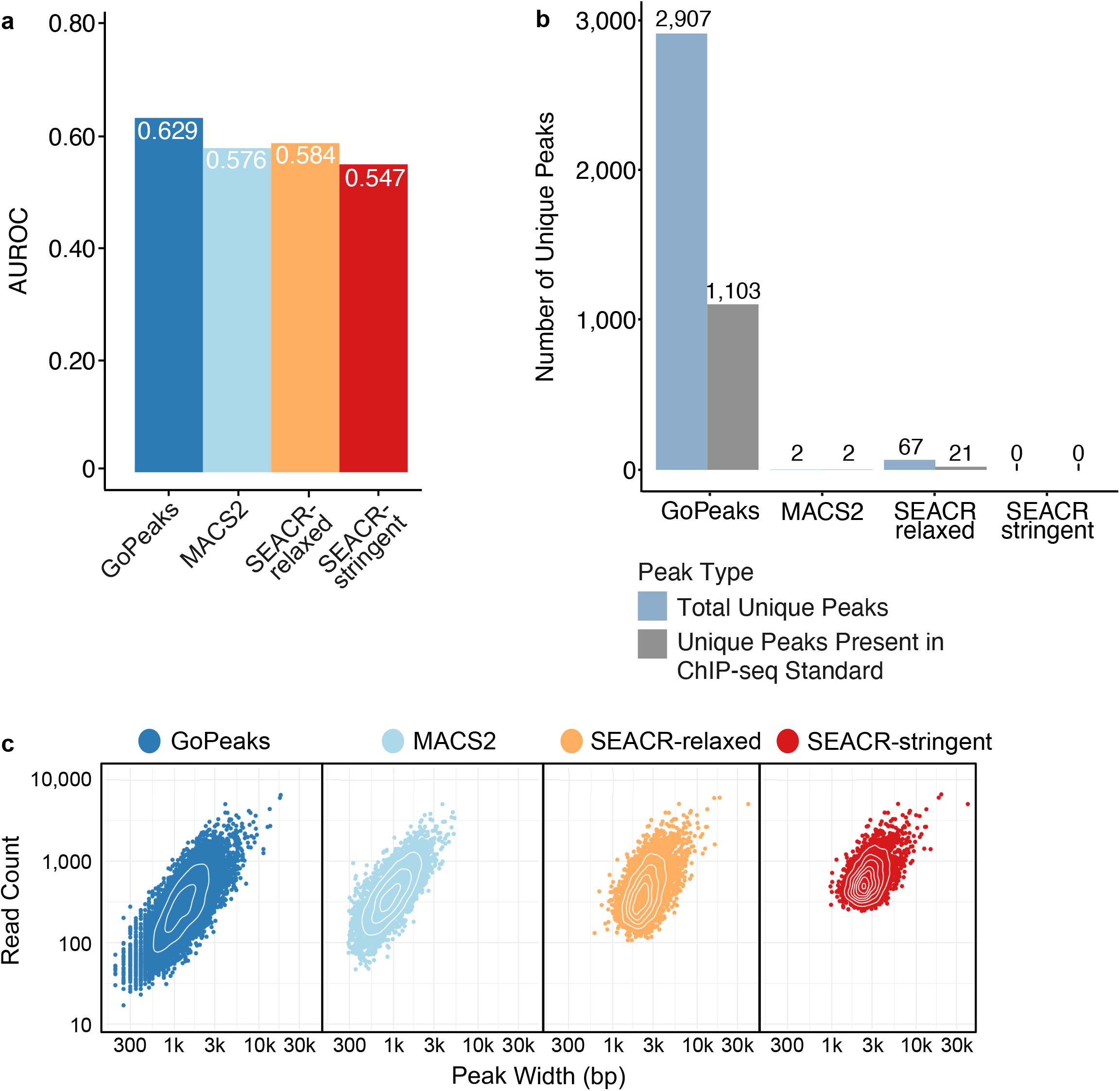
GoPeaks demonstrates improved sensitivity and specificity in identifying H3K27Ac ChIP-seq standard peaks from CUT&Tag data. **a**. Area under the ROC curve (AUROC) for each peak calling method. Each bar is labeled by the AUROC value it represents. Colors indicate the peak calling method. **b**. Comparison of unique peaks identified by each peak calling algorithm and how many are also present in the ChIP-seq standard. Each bar is labeled by the number of peaks it represents. Colors indicate the peak type. **c**. Distribution of read counts by peak width. Each dot represents the read count and peak width of a single detected peak. Colors indicate the peak calling method.

